# Spatially coherent diffusion of human RNA Pol II depends on transcriptional state rather than chromatin motion

**DOI:** 10.1101/2022.01.19.476954

**Authors:** Roman Barth, Haitham A. Shaban

## Abstract

Gene transcription by RNA polymerase II (RNAP II) is a tightly regulated process in the genomic, temporal, and spatial context. Transcriptionally active genes often spatially cluster at RNA Pol II foci, called transcription factories, causing long-range interactions between distal sites of the genome. Recently, we have shown that chromatin exhibits spatially long-range coherently moving regions over the entire nucleus, and transcription enhances this long-range correlated DNA movement. Yet, it remains unclear how the mobility of RNA Pol II molecules is affected by transcription regulation and whether this response depends on the coordinated chromatin movement. We applied our Dense Flow reConstruction and Correlation method to analyze nucleus-wide coherent movements of RNA Pol II in living human cancer cells. We quantify the spatial correlation length of RNA Pol II in the context of DNA motion. We observe a spatially coherent movement of RNA Pol II molecules over ~1 μm, considerably less than for DNA, suggesting that spatially coherent RNA Pol II motion does not solely result from the DNA motion. In contrast to DNA, inducing transcription in quiescent cells decreased the coherent motion of RNA Pol II, while the inhibition of transcription elongation by using DRB slightly increased coherent RNA Pol II motion. The spatially coherent movement of RNA Pol II domains is affected by the transcriptional state and largely independent of the underlying chromatin domains. Our study reveals the nucleus-wide interplay between chromatin and RNA Pol II in the dynamic regulation of chromatin organization.

## Introduction

Genome structure, dynamics, and transcription are highly coordinated to ensure proteins punctually find the proper places on the genome for a correct gene expression [1]. This interplay between dynamics of genome organization and transcription alters and supports the activity of the other [2]. Transcription by RNA Polymerase II (Pol II) takes place for all protein-coding genes in eukaryotic genomes and is vital for many physiological processes [3]. Transcription often takes place in so-called transcription factories, domains of clustered transcription factors, whose formation has been explained by liquid-liquid phase separation [4–6]. These transcription factories are highly dynamic macromolecular that permit transcription initiation and elongation [7,8]. Transcription factories have been proposed to strongly bind DNA, likely regulatory elements, thereby constraining chromatin diffusion nuclear-wide [9,10].

Advances in live-cell imaging and genetic modification tools have revealed the dynamic properties of both RNA Pol II and chromatin and their importance for transcriptional regulation [11,12]. Live-cell imaging of endogenous RNA Pol II using fluorescence recovery after photobleaching uncovered several dynamic states of RNA Pol II [13]. Single-molecule tracking technologies have been applied to image and quantify single RNA Pol II molecules as they bind at non-specific sites throughout the genome [14,15], and a single gene [12,16,17]. Nucleus-wide analysis of RNA Pol II in single living cells has been also analyzed and mapped at high resolution, using a new approach called Hi-D [10]. Recently, super-resolution imaging studies showed the physical relationship between RNA Pol II and chromatin clutches, but the cell fixation hindered deducing dynamic information of RNA Pol II [18,19]. In conclusion, it remains elusive if and how the diffusion of RNA Pol II is coordinated within the context of its surrounding chromatin.

Recently, we developed Dense Flow reConstruction and Correlation (DFCC), a method to study spatial and temporal long-range correlations of abundant nuclear macromolecules over entire single nuclei [20]. DFCC combines light microscopy and computer vision (Optical Flow) technology to reconstruct the dynamics of bulk chromatin in diffraction-limited optical microscopy images at nanoscale resolution throughout the entire nucleus simultaneously. DFCC does not rely on the identification and tracking of single molecules, as compared to single-particle tracking methods, and can thus be applied to abundant nuclear proteins, allowing the estimation of their dynamics in living cells [11,21]. By applying DFCC to genome dynamics during transcription, we detected the formation of long-range correlated chromatin domains, extending up to several micrometers across the nucleus [20,22]. These large coherent domains were reduced by the inhibition of transcription elongation [20].

Here, we apply DFCC to RNA Pol II in non-transcribing and actively transcribing cells to study the nucleus-wide coherent movements of RNA Pol II in living human cells. We find that RNA Pol II exhibits spatially coherent movement, which is markedly reduced upon transcription activation but only partially affected by inhibition of transcription elongation. We then calculate the spatial correlation length of RNA Pol II in the context of DNA motion. In contrast to DNA, inducing transcription in quiescent cells decreased the coherent motion of RNA Pol II. We thus conclude that the spatially coherent movement of RNA Pol II domains is largely independent of the underlying chromatin domains.

## Results

To investigate whether the mobility of RNA Pol II molecules exhibits coherent movement within the nucleus, we applied our recently developed Dense Flow reConstruction and Correlation method (DFCC; Figure 1a) [20]. DFCC applies Optical Flow on time-resolved fluorescence image series to estimate the flow field of fluorescently labeled macromolecules between successive images (Figure 1b). The displacement magnitude and direction are obtained for every pixel across the entire nucleus, which allows computing the spatial and temporal correlation function of both flow magnitude and direction (Figure 1c). A quantitative description of these correlation functions is obtained by regression to the Matérn covariance function (Figure 1c inset; Methods). To characterize flow fields of RNAP II, the correlation length ξ is obtained from the regression, describing how quickly correlations decay over distance [23]. Illustratively, Supplementary Figure S1 displays flow fields exhibiting various degrees of magnitudinal and directional correlation. In brief, the magnitudinal and directional correlation length quantify the distance over which the magnitude (direction, respectively) of displacement vectors is correlated (or “similar”). A flow field may in principle exhibit correlations in neither, either displacement magnitudes or directions or both. Suppose a flow field of a collection of biomolecules locally moving in the same direction, but each with a different speed (or magnitude), the directional correlation length will be large, while the magnitudinal correlation will decay quickly. Thus, the magnitudinal correlation length will be low (e.g., Figure S1b).

**Figure 1:**
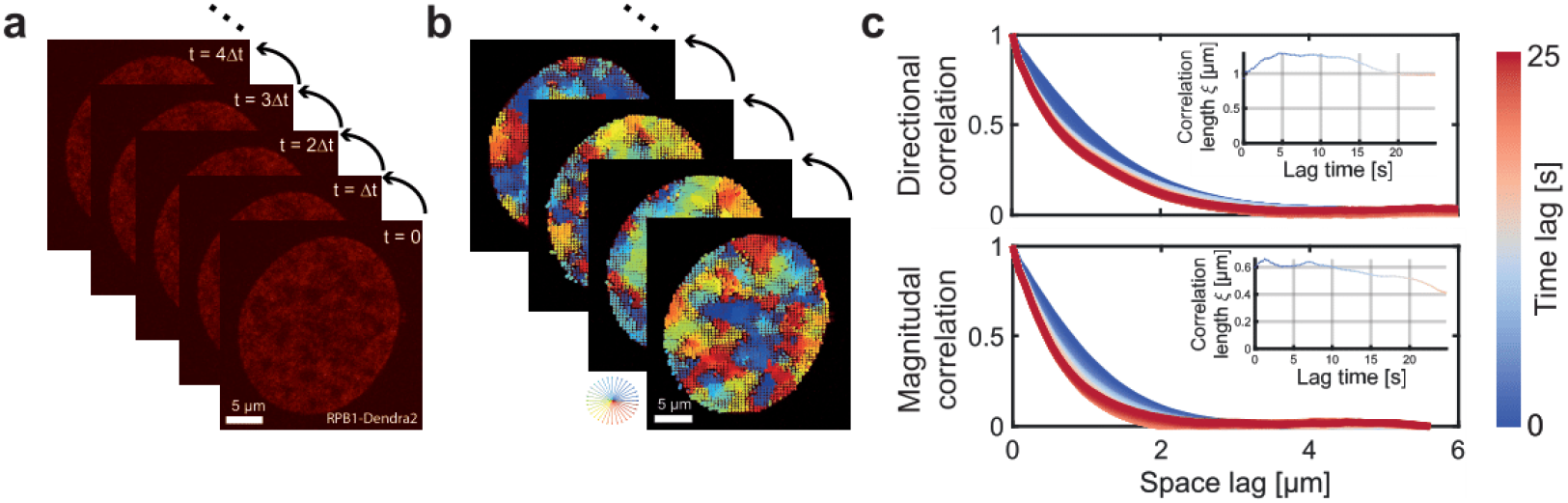
DFCC workflow. **a)** RNAP-Dendra2 stained nuclei of U2OS cells are imaged with a time interval of Δt by confocal microscopy. **b)** Flow fields between successive images are computed using Optical Flow. **c)** The spatial correlation in flow field direction (upper panel) and flow magnitude (lower panel) is computed over increasing space lags (averaged over the two spatial dimensions) and over accessible time lags (from blue to red). The spatial directional and magnitudinal correlation length, respectively, is obtained via regression to the Whittle–Matérn covariance model for every time lag (insets).

Time-resolved movies of RNA Pol II (RPB1-Dendra2, subunit of RNA Pol II) in the U-2 OS human osteosarcoma cell line were recorded at 150 frames with an exposure time of 0.2 s. Cells were grown in a medium in the absence of serum for 24 h to obtain a reference state of inactive transcription [9,10,17,24]. In the absence of transcription, the correlated motion of RNA Pol II molecules was detected in both flow direction and magnitude, while the directional correlation length slightly increased with increasing time lag (Figure 2).

**Figure 2:**
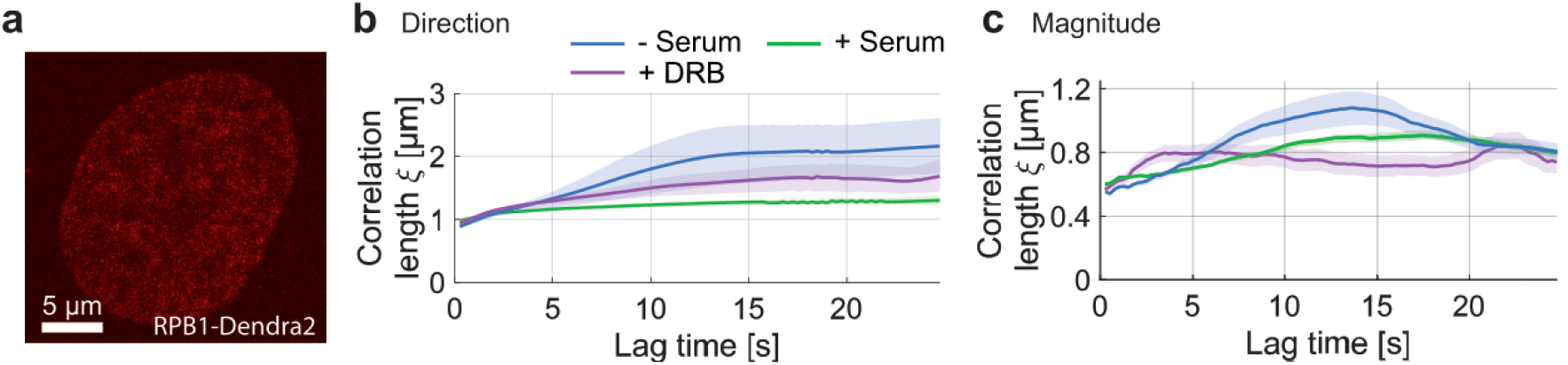
Spatial correlation of RNAP dynamics in the absence of serum, upon serum-stimulation and DRB treatment. **a)** Exemplary RNAP-Dendra2 stained nucleus. **b)** Directional and **c)** magnitudinal correlation length of RNAP over increasing time lag.

To study the correlated motions of RNA Pol II and its changes at different transcriptional stages, we imaged RNA Pol II in two different conditions, serum-starved (inactively transcribing cells) and serum-stimulated cells (actively transcribing). Our and other studies have shown that transcription is largely turned off in serum-starved cells, yet promptly activated when serum is added for 15 minutes to the same cells [9,10,17]. For serum-starved cells, DFCC analysis showed that the correlation length for RNA Pol II is time-dependent with a directional correlation length increasing from ≈ 1 μm to ≈ 2 μm within a lag time of 10 seconds. In contrast, the directional correlation length is time-independent (*ξ* ≈ 1 μm) for serum-stimulated cells as the normal growth condition (Figure *2*b). Note that the serum-stimulated condition showed the same dynamic response as the normal growth condition [9]. The difference in the directional correlation length between coherent domains could be explained by the dynamic clustering of RNA Pol II molecules at transcribed sites due to transcription activation [1,17]. The time-dependence of the magnitudinal correlation length may reflect the dynamic movement of RNA Pol II molecules in forming their coherent domains at their target sites. The correlation of displacement magnitudes in actively transcribing cells was also slightly lower compared to the transcriptionally inactive case for most time lags, yet qualitatively following a similar trend (Figure *2*c The non-vanishing spatial correlation of RNA Pol II molecules within the nucleus might hint at the clustering of RNA Pol II during the formation of transcription factories [25–27].

Similarly, we tested how the spatial correlation of RNA Pol II responds to the inhibition of transcription elongation by using 5,6-Dichloro-1-β-D-ribofuranosylbenzimidazole (DRB). DRB pauses transcription elongation by interrupting cyclin-dependent kinase 9 (CDK9) phosphorylation [28,29]. Several imaging studies of RNA Pol II proposed that transcription inhibition using DRB drug prevents the dissociation of the formed RNA Pol II clusters from the promoter-proximal paused state [14,17,30,31]. DRB was added to cells grown in the normal condition (serum-supplied). In comparison to serum-stimulated cells, DRB treated cells showed a slight increase in directional correlation length and no change in the magnitude of correlation (Figure *2*b, c). Our results suggest that the spatially coherent movement of RNA Pol II domains is affected by transcription initiation and to a smaller extent by elongation.

We then asked whether the coordinated RNA Pol II motion depends on the spatially coherent motion of chromatin [20]. For this aim, the correlation length of RNA Pol II dynamics was compared to the one of chromatin (DNA labeled SiR–Hoechst) in the same cell line and conditions (Figure 3 a, b). Contrary to RNA Pol II, chromatin showed an increase in correlation length (*ξ* ≈ 6 μm) in serum-stimulated cells (active transcription state) and reaches a plateau after a lag time of 20 sec (Figure 3a). Similarly, the correlation length of DNA was reduced in cells treated with DRB (plateau value *ξ* ≈ 4 μm and *ξ* ≈ 0.6 μm for flow direction and magnitude, respectively) (Figure 3a, b). Since chromatin exhibits a considerably longer correlation length than found for RNA Pol II and contrasting responses of the correlation length upon transcriptional activation/inhibition, we conclude that the observed spatial correlation of nuclear RNA Pol II molecules cannot solely be explained by its binding to DNA (see Discussion). Using high-resolution imaging, recent studies also revealed the dissimilarity of RNA Pol II and DNA mobility —characterized by diffusion constant and anomalous exponent— in serum stimulated cells [9,10]. These studies explained the constrained motion of chromatin by the formation of transcription “hubs” or factories, which globally constrains chromatin motion. The short correlation length (*ξ* ≤ 2 μm) of RNA Pol II may reflect the presence of transcription factories, to which freely diffusing RNA Pol II molecules are attracted. In contrast, the correlation length of chromatin in actively transcribing cells is roughly 3-fold higher, hinting to the possibility that clustering of RNA Pol II modulates the spatial organization of chromatin on a scale spanning several micrometers. This finding might be explained by the transcription factor model where specific and nonspecific protein-chromatin interactions create protein and chromatin clusters in the size range of transcriptional condensates (0.1 to 1.0 μm) [32,33]. RNA Pol II clustering, even in the absence of active transcription, therefore affects the spatial chromatin organization and possibly helps to reactivate transcription [34] (see Discussion).

**Figure 3:**
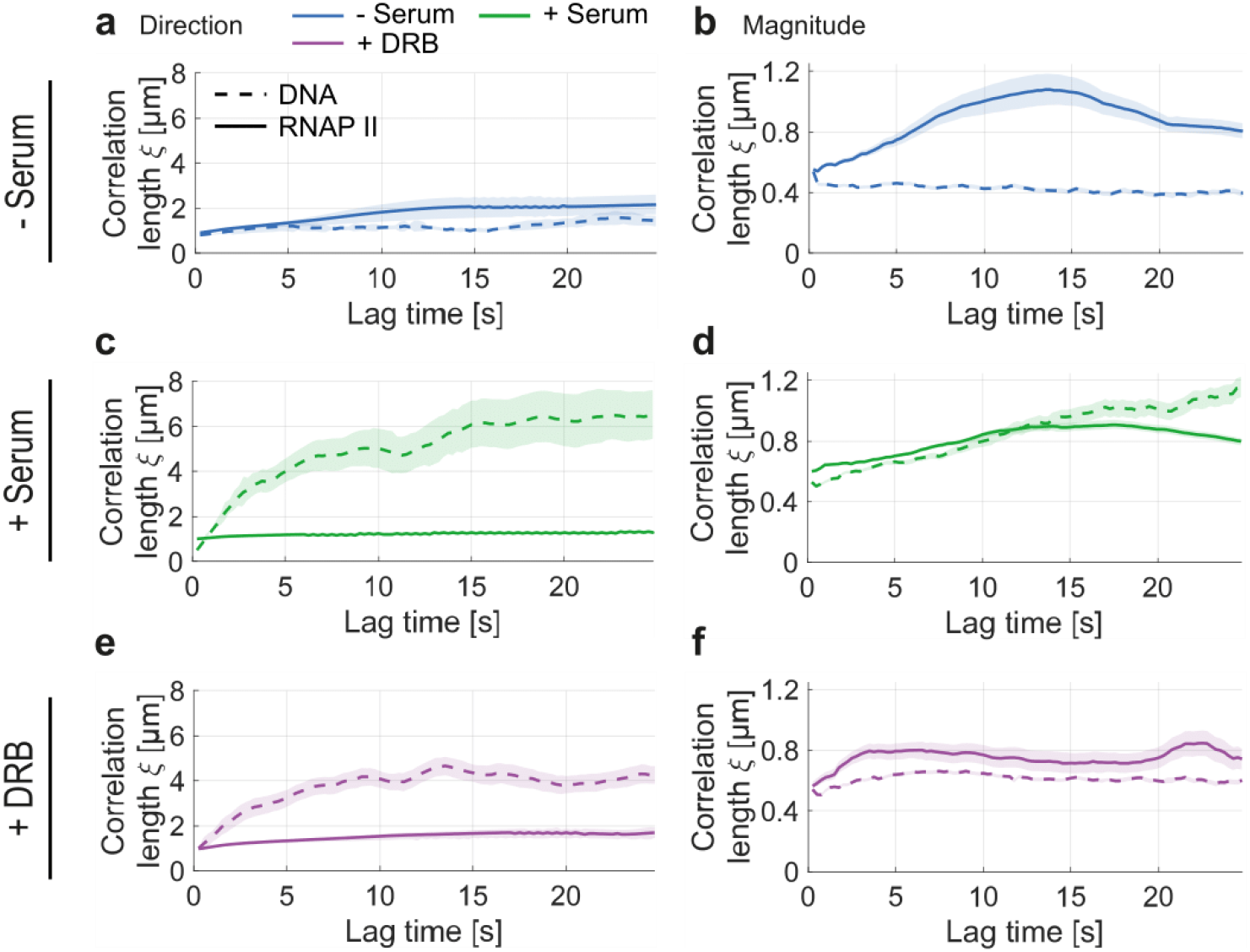
Spatial correlation of DNA and RNA Pol II dynamics for inactive, active, and stalled transcription. **a)** Superimposed directional correlation length for DNA and RNA Pol II in the absence of serum. **b)** Analogous for the magnitudinal correlation. **c-d)** Analogous for serum-stimulated cells. **e-f)** Analogous for DRB treatment in the presence of serum. While DNA dynamics become spatially correlated upon serum stimulation, RNA Pol II’s directional and magnitudinal correlation decreases slightly. Upon stalling RNA Pol II at the initiation step by addition of DRB to the medium, RNA Pol II’s directional correlation slightly increases, while the opposite trend is observed for DNA dynamics. In contrast, DRB treatment reduces the magnitudinal correlation length of both RNA Pol II and DNA.

## Discussion

In this work, we have demonstrated the capability of DFCC for detecting the spatially coordinated movement of RNA Pol II motion at nano-scale resolution. We performed nucleus-wide live-cell imaging of RNA Pol II at different transcriptional states and in the context of chromatin motion. We found that RNA Pol II molecules move in a spatially correlated manner (the correlation length reaching up to ≈ 2 μm), yet its range is considerably reduced compared to the motion of chromatin and exhibits the opposite trend upon serum stimulation and DRB treatment (Figure 4). The formation of these spatially coherent RNA Pol II domains may be explained by the assembly of RNA Pol II into “transcription factories” which were previously observed in fixed cells using super-resolution techniques [25–27]. We report a decrease in the correlation length of RNA Pol II molecules upon transcription activation (Figure 4). This decrease may be due to the organization of unbound molecules into small, clustered regions such as transcription factories [32,35]. Upon inhibiting transcription elongation by DRB treatment, the spatial correlation of RNA Pol II only partially recovers the non-transcribing state, hinting to a combined action of transcription initiation and elongation to the observed spatial coherence of RNA Pol II motion.

**Figure 4:**
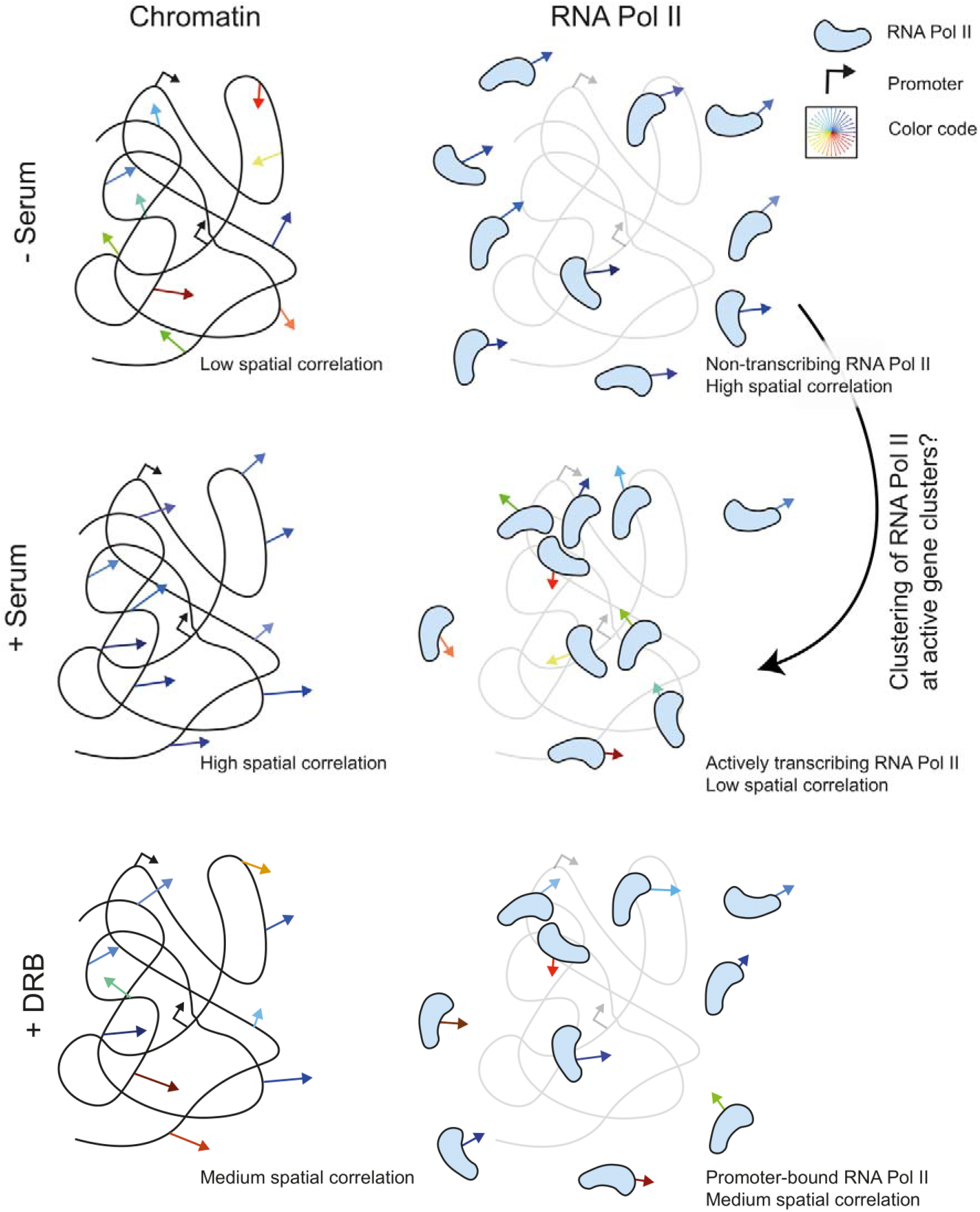
Schematic representation of the observed spatially coherent motion of chromatin versus RNA Pol II in inactive, active, and promoter-paused states of transcription. Spatially coherent chromatin and RNA Pol II motion exhibit opposite trends upon transcription stimulation.

Some plausible scenarios could potentially explain our observations. It is well established that chromatin moves in a spatially coherent manner [20,36,37], while independently diffusing molecules do not exhibit spatially coherent dynamics. Given that RNA Pol II is a DNA-binding protein, the observed spatial correlation may result from the binding of a fraction of RNA Pol II molecules to DNA. Nucleus-wide imaging of RNA Pol II would thus sample both DNA-unbound and DNA-bound fractions. If all RNA Pol II would be DNA-bound, the correlation length of RNA Pol II should recover the one of chromatin and there would be no correlation if no RNA Pol II is bound to DNA. If this argument would be correct and RNA Pol II only exhibits spatially correlated motion when bound to DNA, however, the correlation length of DNA should set an upper limit to the correlated motion of RNA Pol II in all conditions. This is in stark contrast with the fact with the observation that RNA Pol II exhibits a higher correlation length (both in direction and magnitude) than DNA in serum-starved cells (Figure *3*c-d).

A second plausible scenario is that the observed spatial correlation is due to RNA Pol II transcribing chromatin in trains of up to tens of protein copies. Transcription proceeds at an average speed of 2-4 kb/min [13,38,39]. Within the acquisition time of our experiments (30 seconds), we thus expect an average transcribed length of 1-2 kb. Taking the upper limit and stretching such a segment fully, we arrive at a maximum length of the transcribed gene of roughly 2000 *bp* · 0.34 *nm* ≈ 0.7 *μm.* More conservative estimates arrive at an average length of a 10 kb gene around 0.5 *μm* [40]. Most genes are indeed packed with nucleosomes and considerably smaller than the contour length of DNA alone. Given that we observe a directional correlation length ≥ 1 *μm,* we find it highly unlikely that the observed spatial correlation of RNA Pol II is due to the transcription of highly active genes. A molecular picture to how RNA Pol II molecules can exhibit spatially correlated movement and how transcription initiation and elongation separately contribute to this phenomenon is currently lacking and requires further research. We envision that the combination of nucleus-wide live-cell imaging of chromatin and transcription factors, sequencing and modeling approaches will be a powerful approach to answer outstanding questions in how chromatin organization and dynamics interact and shape and are shaped by transcription

## Methods

### Cell culture, treatment, and imaging

Cell culture, starvation, stimulation, treatment, and imaging are performed as described in (18). Briefly, a human U2OS osteosarcoma cell line (for DNA imaging) was maintained in Dulbecco’s modified Eagle’s medium including phenol red-free (Sigma-Aldrich). This cell line stably expresses RPB1 fused with Dendra2 as already described in (13). The medium was supplemented with 10% fetal bovine serum, 1□mM sodium pyruvate (Sigma-Aldrich), Glutamax containing 50□μg/ml gentamicin (Sigma-Aldrich), and G418 0.5□mg/ml (Sigma-Aldrich). Cells were cultivated at 37 °C with 5% CO_2_.

For serum starvation, cells were plated with a serum-free medium and incubated for 24□h at 37 °C before imaging. Before imaging, cells were mounted in the L-15 medium. For stimulation, 10% fetal bovine serum was added to the medium for 15□min. Serum removal arrests cells in G0 and due to the short stimulation with serum, cells are expected to be in the G1 phase of the cell cycle.

For DRB treatment, cells were treated by adding 100□μM DRB (Sigma-Aldrich) to the L-15 imaging medium that contained 10% fetal bovine serum.

Cell fixation. First, the U2OS cells were gently washed with a pre-warmed (37 °C) phosphate-buffered saline (PBS), then cells were incubated in 4% (vol/vol) paraformaldehyde in PBS for 10–20 min at room temperature. Just before imaging, cells were washed with PBS (three times, 5 min each). Images were recorded at room temperature in PBS.

For DNA staining, U2OS cells were labeled by SiR-DNA (SiR-Hoechst) at a final concentration of 2□μM at 37□°C for 30–60 min. Before imaging, the medium was changed to the L-15 medium for live imaging.

Image series of RNA Pol II were recorded as in (9). Image series of 150 frames were acquired at 5 frames per second using a Nipkow-disk confocal system. For Dendra2 excitation, a single wavelength of 488 nm (Coherent) at 10% laser power passed through a 100 ×□oil immersion objective was applied. Images were detected on a cooled electron-multiplying charge-coupled device camera (iXon Ultra 888), with a sample pixel size of 88□nm. The same imaging conditions were applied for DNA imaging, but with an excitation wavelength of 647□nm (Coherent) at 20% laser power.

### Data analysis

Single fluorescent nuclei were manually cropped and then processed as described in [20]. In brief, denoised and drift-corrected images were subject to Optical Flow [41], which results in an estimation of the average displacement of fluorescently stained molecules between consecutive frames for every pixel within the nucleus. The DFCC method first computes the spatial autocorrelation *r*(Δ*x*, Δ*y*) of the resulting flow fields’ magnitude and direction for all accessible lag times between flow fields via

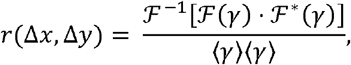

and averages over time. Here, 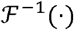 is the inverse Fourier transformation, and 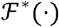 is the complex conjugate of the Fourier transformation. The 2D correlation function was projected as a radial average onto one dimension using the space lag ρ^2^ = Δ*x*^2^ + Δ*y*^2^. Thus, the correlation function turns to a function of the space lag only, i.e., *r* = *r*(*ρ*). Regression of these correlation curves over distance was performed to the Whittle-Matérn covariance function [42]

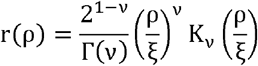

where Γ(·) denotes the gamma function; K_v_(·) the modified Bessel function of the second type of order v, ξ is the correlation length and ν denotes the smoothness parameter. While ξ describes the long-range behavior (over which distance are two measurements correlated), the smoothness parameter ν describes the local, high-frequency component of the correlations (if the flow field can be described as rough or smooth, thus direction/magnitude of spatially close displacement vectors tend to (not) align on a local scale). While ν is also a regression parameter, we found no significant difference in v across the transcriptional conditions and therefore excluded the analyses of the smoothness parameter.

We validated that all obtained correlation lengths are above the detection limit of DFCC by applying DFCC to the image series of chemically fixed cells (Supplementary Figure S2). We found that the obtained correlation lengths both for chromatin and RPB1-Dendra2 are overall well below the values found for living cells and do not show any dependence on the time lag, thus setting the sensitivity baseline for the DFCC method for the live-cell imaging of chromatin and RNA Pol II, respectively.

Error bars of the shown correlation lengths display the standard error of the mean (SEM) of 19, 21, and 21 cells for serum-starved, serum-stimulated, and DRB-treated cells, respectively. For fixed cell analyses, 17 and 22 cells were analyzed for chromatin and RNA Pol II-stained nuclei.

### Generation of spatially correlated flow fields

Flow fields in Figure S1 were generated on a 100 x 100 grid by separately sampling the displacement vectors from a distribution according to the simulated scenario. For flow fields without correlation (in either magnitude or direction or both), the magnitude was sampled from a normal distribution with a mean of 0.5 and a standard deviation of 0.05. The direction values were sampled from a uniform distribution from 0 to 2*π*. Spatially correlated values (for either magnitude or direction or both) were generated as a stochastic two-dimensional multi-fractal random field [43].

## Acknowledgments

We acknowledge support from the Pôle Scientifique de Modélisation Numérique, ENS de Lyon for providing computational resources.

## Author contributions

H.A.S. designed and supervised the project. R.B. performed data analysis. H.A.S. carried out experimental work. H.A.S. and R.B. interpreted the results. H.A.S. and R.B. wrote the manuscript.

## Data and materials availability

All data needed to evaluate the conclusions in the paper are present in the paper. Additional data related to this paper may be requested from the authors.

## References

1. Shaban HA, Barth R, Bystricky K. Navigating the crowd: visualizing coordination between genome dynamics, structure, and transcription. Genome Biol. 2020.

2. van Steensel B, Furlong EEM. The role of transcription in shaping the spatial organization of the genome. Nat. Rev. Mol. Cell Biol. 2019.

3. Schier AC, Taatjes DJ. Structure and mechanism of the RNA polymerase II transcription machinery. Genes Dev. 2020.

4. Sabari BR, Dall’Agnese A, Boija A, Klein IA, Coffey EL, Shrinivas K, et al. Coactivator condensation at super-enhancers links phase separation and gene control. Science (80-). 2018;

5. Cho WK, Spille JH, Hecht M, Lee C, Li C, Grube V, et al. Mediator and RNA polymerase II clusters associate in transcription-dependent condensates. Science (80-). 2018;361:412–5.

6. Pancholi A, Klingberg T, Zhang W, Prizak R, Mamontova I, Noa A, et al. RNA polymerase II clusters form in line with surface condensation on regulatory chromatin. Mol Syst Biol. 2021;

7. Ghamari A, van de Corput MËPC, Thongjuea S, Van Cappellen WA, Van Ijcken W, Van Haren J, et al. In vivo live imaging of RNA polymerase II transcription factories in primary cells. Genes Dev. 2013;

8. Rippe K, Papantonis A. RNA polymerase II transcription compartments: from multivalent chromatin binding to liquid droplet formation? Nat. Rev. Mol. Cell Biol. 2021.

9. Nagashima R, Hibino K, Ashwin SS, Babokhov M, Fujishiro S, Imai R, et al. Single nucleosome imaging reveals loose genome chromatin networks via active RNA polymerase II. J Cell Biol. 2019;

10. Shaban HA, Barth R, Recoules L, Bystricky K. Hi-D: nanoscale mapping of nuclear dynamics in single living cells. Genome Biol. 2020;21:95.

11. Shaban HA, Seeber A. Monitoring the spatio-temporal organization and dynamics of the genome. Nucleic Acids Res. 2020.

12. Li J, Dong A, Saydaminova K, Chang H, Wang G, Ochiai H, et al. Single-Molecule Nanoscopy Elucidates RNA Polymerase II Transcription at Single Genes in Live Cells. Cell. 2019;

13. Steurer B, Janssens RC, Geverts B, Geijer ME, Wienholz F, Theil AF, et al. Live-cell analysis of endogenous GFP-RPB1 uncovers rapid turnover of initiating and promoter-paused RNA Polymerase II. Proc Natl Acad Sci. 2018;

14. Cisse II, Izeddin I, Causse SZ, Boudarene L, Senecal A, Muresan L, et al. Real-time dynamics of RNA polymerase II clustering in live human cells. Science (80-). 2013;341:664–7.

15. Uchino S, Ito Y, Sato Y, Handa T, Ohkawa Y, Tokunaga M, et al. Live imaging of transcription sites using an elongating RNA polymerase II–specific probe. J Cell Biol [Internet]. 2021;221:e202104134. Available from: https://doi.org/10.1083/jcb.202104134

16. Forero-Quintero LS, Raymond W, Handa T, Saxton MN, Morisaki T, Kimura H, et al. Live-cell imaging reveals the spatiotemporal organization of endogenous RNA polymerase II phosphorylation at a single gene. Nat Commun. 2021;

17. Cho WK, Jayanth N, English BP, Inoue T, Andrews JO, Conway W, et al. RNA Polymerase II cluster dynamics predict mRNA output in living cells. Elife. 2016;5.

18. Castells-Garcia A, Ed-daoui I, González-Almela E, Vicario C, Ottestrom J, Lakadamyali M, et al. Super resolution microscopy reveals how elongating RNA polymerase II and nascent RNA interact with nucleosome clutches. Nucleic Acids Res [Internet]. 2022;50:175–90. Available from: https://doi.org/10.1093/nar/gkab1215

19. Miron E, Oldenkamp R, Brown JM, Pinto DMS, Xu CS, Faria AR, et al. Chromatin arranges in chains of mesoscale domains with nanoscale functional topography independent of cohesin. Sci Adv. 2020;

20. Shaban HA, Barth R, Bystricky K. Formation of correlated chromatin domains at nanoscale dynamic resolution during transcription. Nucleic Acids Res. 2018;46.

21. Agbleke AA, Amitai A, Buenrostro JD, Chakrabarti A, Chu L, Hansen AS, et al. Advances in Chromatin and Chromosome Research: Perspectives from Multiple Fields. Mol Cell. 2020;

22. Barth R, Bystricky K, Shaban HA. Coupling chromatin structure and dynamics by live superresolution imaging. Sci Adv [Internet]. American Association for the Advancement of Science; 2020;6. Available from: https://advances.sciencemag.org/content/6/27/eaaz2196

23. Stein ML. Interpolation of spatial data: some theory for kriging. Springer; 1999.

24. Ray S, Panova T, Miller G, Volkov A, Porter ACG, Russell J, et al. Topoisomerase IIα promotes activation of RNA polymerase I transcription by facilitating pre-initiation complex formation. Nat Commun [Internet]. 2013;4:1598. Available from: http://www.nature.com/doifinder/10.1038/ncomms2599

25. Carter DRF, Eskiw C, Cook PR. Transcription factories. Biochem. Soc. Trans. 2008.

26. Rieder D, Trajanoski Z, McNally JG. Transcription factories. Front Genet. 2012;

27. Papantonis A, Cook PR. Transcription factories: Genome organization and gene regulation. Chem. Rev. 2013.

28. Sehgal PB, Darnell JE, Tamm I. The inhibition of DRB (5,6-dichloro-1-ß-d-ribofuranosylbenzimidazole) of hnRNA and mRNA production in HeLa cells. Cell. 1976;

29. Bensaude O. Inhibiting eukaryotic transcription: Which compound to choose? How to evaluate its activity? Transcription. 2011;

30. Mitchell JA, Fraser P. Transcription factories are nuclear subcompartments that remain in the absence of transcription. Genes Dev. 2008;

31. Cho WK, Jayanth N, Mullen S, Tan TH, Jung YJ, Cissé II. Super-resolution imaging of fluorescently labeled, endogenous RNA Polymerase II in living cells with CRISPR/Cas9-mediated gene editing. Sci Rep. 2016

32. Brackley CA, Taylor S, Papantonis A, Cook PR, Marenduzzo D. Nonspecific bridging-induced attraction drives clustering of DNA-binding proteins and genome organization. Proc Natl Acad Sci [Internet]. 2013;110:E3605–11. Available from: http://www.pnas.org/cgi/doi/10.1073/pnas.1302950110

33. Brackley CA, Liebchen B, Michieletto D, Mouvet F, Cook PR, Marenduzzo D. Ephemeral Protein Binding to DNA Shapes Stable Nuclear Bodies and Chromatin Domains. Biophys J. 2017;

34. Zhang S, Übelmesser N, Josipovic N, Forte G, Slotman JA, Chiang M, et al. RNA polymerase II is required for spatial chromatin reorganization following exit from mitosis. Sci Adv. 2021;

35. Chiang M, Brackley CA, Marenduzzo D, Gilbert N. Predicting genome organisation and function with mechanistic modelling. Trends Genet. 2022.

36. Di Pierro M, Potoyan DA, Wolynes PG, Onuchic JN. Anomalous diffusion, spatial coherence, and viscoelasticity from the energy landscape of human chromosomes. Proc Natl Acad Sci [Internet]. 2018;115:7753–8. Available from: http://www.pnas.org/lookup/doi/10.1073/pnas.1806297115

37. Salari H, Di Stefano M, Jost D. Spatial organization of chromosomes leads to heterogeneous chromatin motion and drives the liquid-or gel-like dynamical behavior of chromatin. Genome Res. 2022;32:28–43.

38. Darzacq X, Shav-Tal Y, de Turris V, Brody Y, Shenoy SM, Phair RD, et al. In vivo dynamics of RNA polymerase II transcription. Nat Struct Mol Biol. 2007;14:796–806.

39. Wada Y, Ohta Y, Xu M, Tsutsumi S, Minami T, Inoue K, et al. A wave of nascent transcription on activated human genes. Proc Natl Acad Sci. 2009;106:18357–61.

40. Leidescher S, Ribisel J, Ullrich S, Feodorova Y, Hildebrand E, Galitsyna A, et al. Spatial organization of transcribed eukaryotic genes. Nat Cell Biol. 2022;24:327–39.

41. Sun D, Roth S, Black MJ. A quantitative analysis of current practices in optical flow estimation and the principles behind them. Int J Comput Vis. 2014;106:115–37.

42. Matérn B. Spatial variation. Springer-Verlag; 1986.

43. Schertzer D, Lovejoy S. Nonlinear Variability in Geophysics: Multifractal Simulations and Analysis. Fractals’ Phys Orig Prop. Boston, MA: Springer US; 1989. p. 49–79.

