## Supplementary Information for "Spatially coherent diffusion of human RNA Pol II depends on transcriptional state rather than chromatin motion"

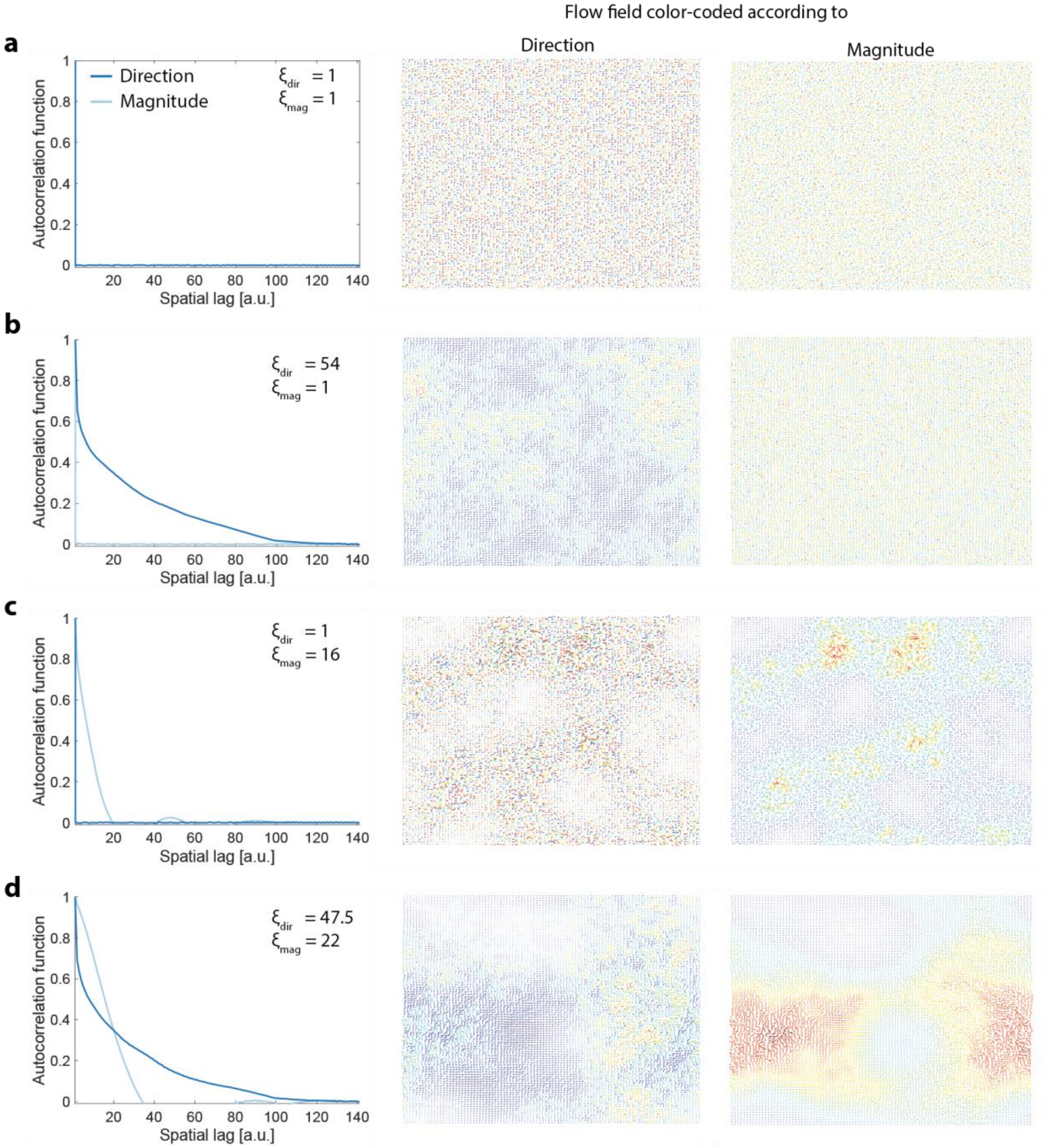


**Figure S1**: Illustrative behavior of the magnitudal and directional correlation length ($\xi_{mag}$ and $\xi_{dir}$, respectively) for four artificial flow fields. The magnitudal and directional autocorrelation length is plotted over spatial lags on the left. The flow fields are colored according to the direction (middle column) and magnitude (right column). **a** Flow field without correlation, **b** with only directional, **c** magnitudal correlation, and **d** both.

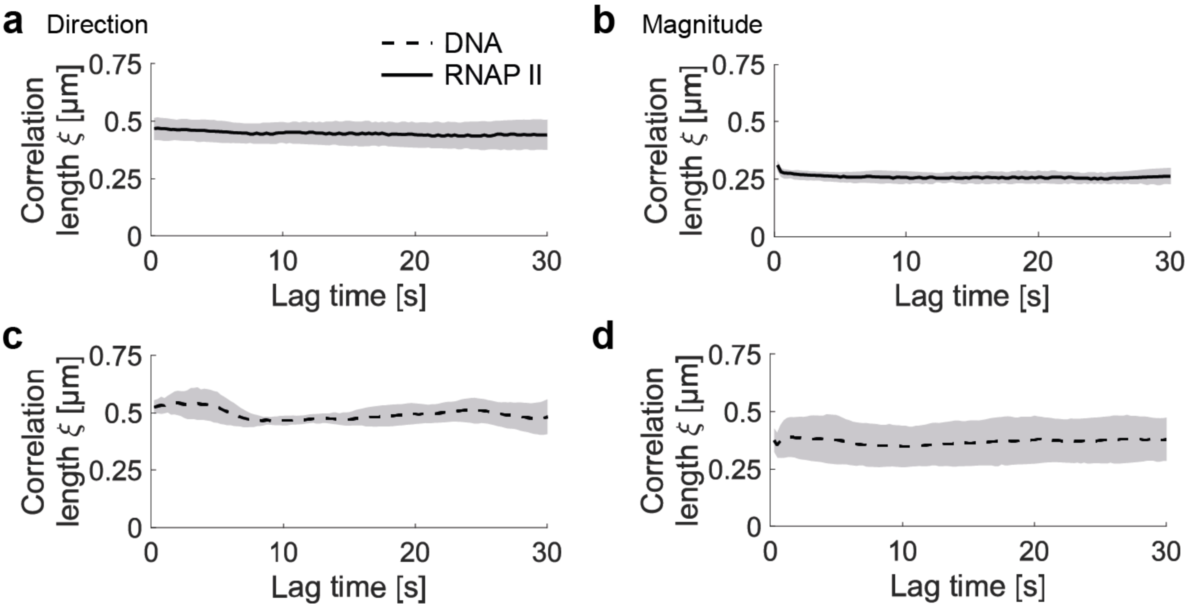


**Figure S2**: Correlation lengths for chemically fixed U2OS cells stained for DNA (SiR-DNA) and RNA Pol II (Rpb1-Dendra2). All obtained correlation lengths are below the values found for living cells and do not depend on the time lag, setting the sensitivity standard for DNA- and RNA Pol II stained nuclei. **a-b** Directional and magnitudinal correlation length for RNA Pol II and **c-d** for DNA.
